# Time dependent stress relaxation and recovery in mechanically strained 3D microtissues

**DOI:** 10.1101/2020.01.25.916874

**Authors:** Matthew Walker, Michel Godin, James L. Harden, Andrew E. Pelling

**Affiliations:** Department of Biology, Gendron Hall, 30 Marie Curie, University of Ottawa, Ottawa, ON, K1N5N5 Canada; Department of Physics, 150 Louis Pasteur pvt., STEM Complex, University of Ottawa, Ottawa, ON K1N 6N5 Canada; Department of Mechanical Engineering, Colonel By Hall, 161 Louis Pasteur, University of Ottawa, Ottawa, ON K1N6N5 Canada; Ottawa-Carleton Institute for Biomedical Engineering, Colonel By Hall, 161 Louis Pasteur, University of Ottawa, Ottawa, ON K1N6N5 Canada; Ottawa Institute of Systems Biology, University of Ottawa, Ontario K1H 8M5, Canada; Institute for Science Society and Policy, Simard Hall, 60 University, University of Ottawa, Ottawa, ON, K1N5N5 Canada; SymbioticA, School of Human Sciences, University of Western Australia, Perth, WA, 6009 Australia

**Author notes:** Author for correspondence Andrew E. Pelling, 150 Louis Pasteur pvt., University of Ottawa, Ottawa, ON K1N 6N5, Canada, Web: http://www.pellinglab.net.

**Keywords:** Microtissue, Viscoelasticity, Cell mechanics, 3D cell culture, Microfabrication

## Abstract

Characterizing the time-dependent mechanical properties of cells is not only necessary to determine how they deform, but also to fully understand how external forces trigger biochemical-signaling cascades to govern their behavior. Presently mechanical properties are largely assessed by applying local shear or compressive forces on single cells in isolation grown on non-physiological 2D surfaces. In comparison, we developed the microfabricated vacuum actuated stretcher to measure tensile loading of 3D multicellular ‘microtissue’ cultures. With this approach, we assessed here the time-dependent stress relaxation and recovery responses of microtissues, and quantified the spatial remodeling that follows step length changes. Unlike previous results, stress relaxation and recovery in microtissues measured over a range of step amplitudes and pharmacological treatments followed a stretched exponential behavior describing a broad distribution of inter-related timescales. Furthermore, despite a performed compendium of experiments, all responses led to a single linear relationship between the residual elasticity and degree of stress relaxation, suggesting that these mechanical properties are coupled through interactions between structural elements and the association of cells with their matrix. Lastly, although stress relaxation could be quantitatively and spatially linked to recovery, they differed greatly in their dynamics; while stress recovery behaved as a linear process, relaxation time constants changed with an inverse power law with step size. This assessment of microtissues offers insights into how the collective behavior of cells in a 3D collagen matrix generate the dynamic mechanical properties of tissues, which is necessary to understanding how cells deform and sense mechanical forces in the body.

## Introduction

The requirements of crawling, dividing and contracting demand that the cell’s cytoskeletal network of structural and motor proteins be tremendously dynamic. This behavior is unique from other soft materials, and gives cells and tissues their distinct elastic and dissipative properties. Defining these properties is not only necessary for understanding how cells deform, but also how cells sense and transduce external mechanical forces into biochemical signals that direct their behavior in the body. In that regard, when cells are stretched or come in contact with a stiffer matrix, there are time-scale dependent conformational changes to adhesion and cytoskeletal proteins, which in turn alter ligand-receptor binding affinities to trigger biochemical signaling cascades ^1,2^. Through this regulation of biochemical signaling, mechanical forces have been linked to normal development and function, as well as disease progression, including bone, muscle, heart and lung disorders and cancer ^3,4^.

In regards to defining the mechanical behavior of living matter, it has long been recognized that cells and tissues exhibit both solid-like elastic and fluid-like viscous properties ^5,6^. Traditionally this behavior, called viscoelasticity, has been described using a network of elastic springs, and viscous dashpots. In particular, when these elements are connected in series, solving the constitutive equations for a step change in length gives an exponential decay in stress with a characteristic time constant(s) that depends upon the elastic modulus of the spring(s) and viscosity of the dashpot(s) ^7^.

With these spring-dashpot models in mind, many studies set out to characterize the viscoelastic behavior of single cells and to link them to specific occurring processes. Although early experimental data could be fit with a single time constant ^8–11^, as the resolution of techniques improved, a power law behavior emerged ^12^. In that regard, the frequency, creep and stress relaxation responses of isolated cells have all been shown to be accurately captured by a single power exponent describing a continuous, featureless distribution of timescales ^12–16^. Further universal observations of power law rheology across different cell types, techniques and following a range of cytoskeletal drugs, has since given traction to the hypothesis that cells belong to a class materials called soft glasses ^12,13^. One notable exception to this featureless rheological behavior seems to occur at short timescales (<1 sec) and under large volumetric deformations Under these conditions, a characteristic behavior has been reported arising from poroelastic effects caused by the redistribution of cytosolic fluid ^17^.

In regard to tissue-level mechanics, while a featureless relaxation behavior has also been reported^6,18^, the field has not reached a consensus on the use of power laws in describing their viscoelastic response. Rather, spring-dashpot models with characteristic timescales remain prominently reported in soft tissue mechanics^19,20^. For instance, the viscoelastic behavior of muscle tissue, particularly when the mechanical response is dominated by actin and myosin kinetics, has been shown to deviate from a power-law behavior, and instead followed a broad distribution of timescales around a characteristic time constant set by acto-myosin activity ^21^. Furthermore, growing cells on a 2D substrate, as largely required for assessing individual cell mechanics, forces an un-natural apical-basal polarity of adhesion complexes. This in turn is known to cause vast differences in the distribution and structure of the cytoskeleton ^22^. Although it remains unclear, it is not unreasonable to suspect the these fundamental changes to cytoskeleton, caused by the dimensionality of the cell’s environment, may alter the mechanical behavior of cells in 3D tissues compared to when studied on 2D substrates. Lastly, characteristic detachment rates of cellular adhesions to matrix proteins may further differentiate tissue-level viscoelastic behavior from the power-law rheological model widely seen in isolated cells ^23^. Therefore although our knowledge of the mechanical behavior of isolated cells has greatly advanced, we lack a complete understanding of how a heterogeneous population of cells within a 3D extracellular matrix establishes the time-scale varying and non-linear properties of tissues. To answer this research question, 3D cell culture methods that enable the assessment of tensile forces have become a keen interest in the field of cell mechanics ^24^. In that regard, both the frequency and stress relaxation response of cells within bulk reconstituted collagen gels have previously displayed a characteristic timescale behavior following standard linear spring-dashpot models ^25–27^. Due to their centimeter-scale, however, these cultures tend to have poor cellular organization and low cell density, suffer from slow experimental throughput, are hard to image, and possess a high diffusive barrier for nutrients. These limitations of bulk 3D cell cultures can be largely overcome by shrinking the cell culture size by adopting a lab-on-chip approach, as in the microtissue model ^28^. In that regard, the high-throughput array of sub-millimeter-scale structures that form around pairs of flexible cantilevers in the microtissue model possess comparable cell alignment and density to human dense connective and muscle tissues, as well as enable high-resolution live-cell imaging and assessment of acto-myosin dynamics ^28–31^. Although quasi-static mechanics (i.e. contractility and stiffness) of microtissues have already received much attention ^28,29,32,33^, the time-dependencies in these 3D cell cultures remain unclear.

Accordingly, in this article we investigated the viscoelastic stress relaxation of microtissue cultures using a microfabricated device that we have developed called the Microtissue Vacuum Actuated Stretcher (MVAS) ^30,31^. The MVAS allows large deformations, simultaneous measurements of tension and live imaging of microtissue cultures. Importantly, the description of the dynamic behavior of 3D microtissue cultures that follows in this article qualitatively differs from previous measurements on isolated cells in 2D culture, and thus, raises important questions on our understanding of how an aggregate population of cells produces the time-dependent and nonlinear mechanical properties of tissues.

## Methods

### MVAS device

MVAS devices were fabricated out of polydimethylsiloxane (PDMS) using mold replication and plasma bonding steps outlined previously ^30,31^. Briefly devices consists of 60 microtissue wells; each containing two cantilevers spaced apart by 500μm and bordered on one side by a vacuum chamber (Fig. 1a). Devices consist of three layers replicated from masters created with standard photolithographic techniques. First, the top device layer forms the open-top microtissue wells and has enclosed vacuum chambers. Second, in the middle is a thin flexible membrane that is fabricated with the cantilevers. Lastly, the bottom layer has enclosed vacuum chambers that match the geometry of the top layer and empty chambers that allow for the middle membrane to stretched in plane. When a vacuum is applied via an external electronic vacuum regulator (SMC ITV0010) controlled through Labview software., the cantilever closest to the vacuum chamber moves to stretch the microtissue. The opposing cantilever is used as a passive force sensor by optically tracking its deflection and utilizing its known spring constant (k_cantilever_=0.834N/m). The dimensions of the force sensing cantilever were empirically chosen to be stiff enough to reduce the amount of creep that inherently occurs during stress relaxation because of our method of force measurement, yet also soft enough in order to have a sufficient signal to noise ratio in tension measurements.

**Fig. 1:**
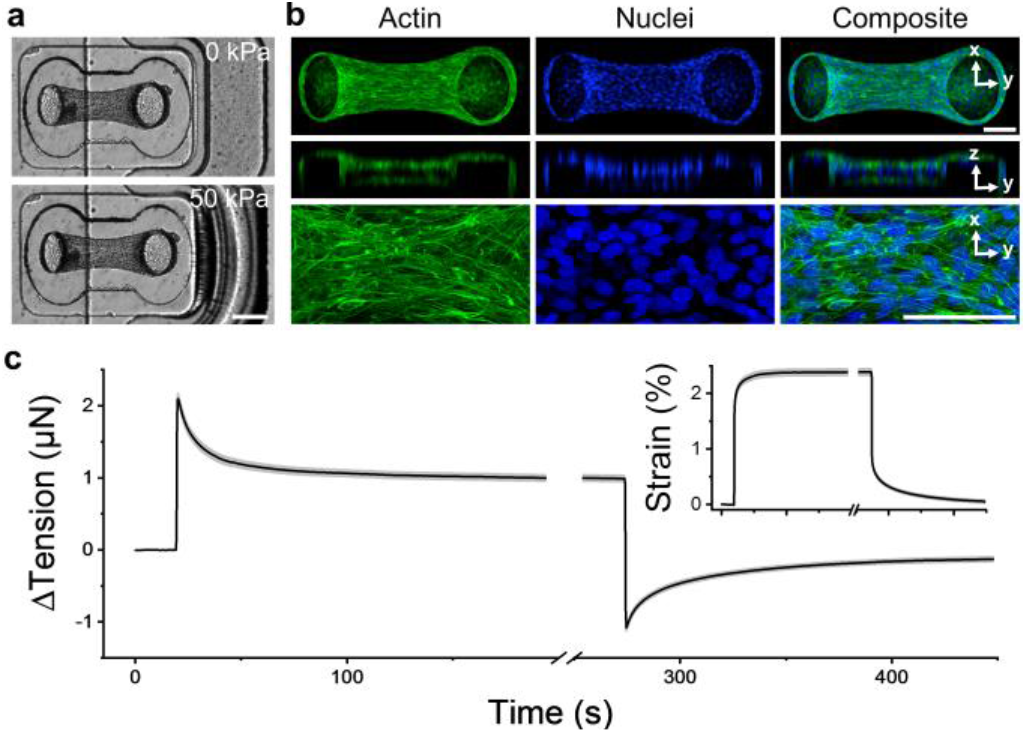
Microtissue tension dynamically relaxed and recovered with changes in length. Microtissues were grown in our MVAS device (a). In the MVAS, changes to microtissue length are driven by applying a regulated vacuum pressure to a chamber that borders one side of each microtissue well. The cantilever closest to the vacuum chamber (shown here on the right) is actuated to stretch the microtissue in plane while changes in tension can then be measured by tracking the deflection of the opposing cantilever (on the left). Microtissues are organized 3D cell cultures freely suspended between the cantilevers. Max projections of confocal stacks, orthogonal views and high magnification images are shown in (b). Both the actin cytoskeleton (green), and the nuclei (blue) are predominately aligned between the cantilevers. Scale bars in (a) and (b) represent 200μm and 100μm, respectively. Following a step in strain (insert), microtissue tension (N=79) sharply increased and then relaxed to a new set point as expected in a viscoelastic solid (c). Then upon returning the microtissue to its original length, the tension recovered.

### Cell culture

NIH3T3 fibroblast (ATCC) cells were cultured in Dulbecco’s Modified Eagle’s Medium (DMEM) (Hyclone Laboratories Inc.) supplemented with 10% fetal bovine serum (FBS), 50mg/ml streptomycin and 50U/ml penicillin antibiotics (all from Hyclone Laboratories Inc.). Cells were grown at 37°C with 5% CO_2_ on 100mm tissue culture dishes (Fisher) until 80-90% confluent.

### Microtissue fabrication

Microtissues consisting of 3T3 fibroblasts in a 3D collagen matrix were cultured as previously described ^28,30^. Briefly, the MVAS was sterilized with 70% ethanol, and treated with 0.2% Pluronic F-127 (P6866, Invitrogen) for two minutes to reduce cell adhesion. 250,000 cells were resuspended in 1.5mg/ml rat tail collagen type I (354249, Corning) solution containing 1x DMEM (SH30003.02, Hyclone), 44 mM NaHCO_3_, 15 mM d-ribose (R9629, Sigma Aldrich), 1% FBS and 1 M NaOH to achieve a final pH of 7.0-7.4. The cell-collagen solution was pipetted into the MVAS and centrifuged to load ~650 cells into each well. The excess collagen was removed and the device was transferred into the incubator for 15 minutes to initiate collagen polymerization. An additional ~130 cells were centrifuged into each well and allowed to adhere to the top of the tissues. Excess cells were washed off. Cell culture media was added and changed every 24 hours.

### Imaging

All images were acquired on a TiE A1-R laser scanning confocal microscope (LSCM) (Nikon) with standard LSCM configurations using appropriate laser lines and filter blocks. To assess morphology, microtissues were fixed *in situ* with paraformaldehyde for 10 minutes and permeabilized with 0.5% Triton-X for 3 minutes. The actin cytoskeleton was stained with Alexa Fluor 546 Phalloidin (Fisher, A22283) and the nuclei were stained with DAPI (Fisher, D1306).

### Force measurements

After three days of static culture, we measured both stress relaxation of microtissues following a step strain and stress recovery after subsequently returning them to their initial length. All measurements were completed at 37°C with 5% CO_2_. Changes in microtissue tension were deduced from the visible deflection of the force-sensing cantilever, which was calculated by subtracting the difference in top and bottom positions. Bright field images of the tops of the cantilevers were captured at 15 frames per second during both stress relaxation and recovery. From these images, the positions of the top of the cantilevers were tracked using pattern matching with adaptive template learning in Labview. To aid in tracking the bottom positions of the force-sensing cantilevers, they were fluorescently labeled by doping the PDMS mixture with Rhodamine-B prior to curing. Fluorescent images taken of the bottom of the cantilevers were captured before and after stress relaxation and recovery experiments. The bottom position of the force-sensing cantilever was measured from these images with a simple centroid algorithm in Matlab. A validation of our approach with an elastic standard and characterization of the noise floor without an attached load are in SI 1.

To investigate nonlinearities in the viscoelastic behavior of microtissues, stress relaxation and recovery were measured for various step strain values. The strain, ε, was defined as the percent change in the length between the innermost edges of the tops of the cantilevers once the microtissue had fully relaxed or recovered (equation 1).

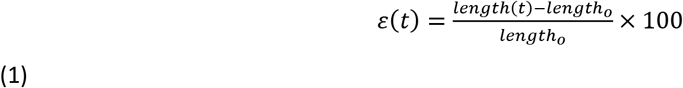

To assess the role of individual cytoskeletal proteins in contributing to the viscoelasticity of microtissues, measurements were taken following 20 minute incubations with either 10μM nocodazole (Noco), a microtubule polymerization inhibitor, 5μM blebbistatin (Bleb), a myosin-II inhibitor, 10 μM cytochalasin D (CytoD), an actin polymerization inhibitor, or 0.5% Triton-X, to decellularize microtissues. 0.5% DMSO was used a loading control. As mechanical properties varied between microtissues, each microtissue was compared to its own pre-treatment value where indicated. To prevent crossover in response from multiple drugs, only a single treatment was administrated to any given microtissue.

### Quantification of microtissue remodeling

To quantify the spatial distribution of remodeling that occurs following a change in length, local strains were estimated across microtissues using a method described previously ^30^. Briefly, starting at the frame immediately following the step change in length, inter-frame displacements were estimated every one second for 100 seconds in Labview at two-pixel spacing across a region of interest using a four level-pyramid based Lucas and Kanade algorithm^34^ with sub-pixel precision and a window size of 17×17 pixels. In Matlab, the displacement field was first smoothed with a LOWESS surface-fitting algorithm, then the inter-frame strain tensor was calculated by finding the gradient of the smoothed displacement field, and finally the inter-frame strain tensor was integrated to estimate the local total strain field. Importantly because the strain distribution was integrated starting the frame immediately following the step change in length, it reflected the viscoelastic behavior of the microtissues rather than how they deform when stretched. Thereby this metric is a measure of the spatial viscoelastic remodeling response.

### Data analysis and statistics

All numerical data are presented as mean ± standard error. Statistical tests as described in the results were performed using Originlab 8.5 (Northampton, MA), with p<0.05 considered statistically significant. Fits to stress relaxation data were performed using Matlab’s curve fitting toolbox.

### Ethics approval

Ethic approval was not required for use of NIH3T3 cell line or for any experiment conducted within this study.

## Results

### Microtissues are viscoelastic

After three days of static culture in the MVAS device, fibroblasts cells had compacted the collagen matrix to form dense 3D microtissues suspended between the cantilevers (fig. 1). Based upon the deflection of the force-sensing cantilever, microtissues had developed a resting tension of 8.7±0.4μN (N=79). The cells were mostly aligned with the direction of tension development shown by both the longitudinally oriented actin cytoskeleton and nuclei (fig. 1b). As been previously reported ^28,29^, these observations indicated that microtissues formed under tensile constraints, as in the MVAS device, are tightly compacted highly organized structures, and in these respects, broadly resemble dense connective and muscle tissues.

In addition to static tensile measurements, the MVAS enables the assessment of mechanical behavior in response to changes in strain through a vacuum-driven planar stretch (fig. 1a). In this regard, following a step change in length, microtissue tension rose sharply and then relaxed to a new equilibrium point as commonly seen in viscoelastic solids (fig. 1c). Upon returning the microtissues to their initial length, the tension dropped past and then slowly recovered to the original resting value (P>0.05, repeated measures t-tests). The stress relaxation and recovery responses were well conversed amongst microtissues (average standard deviation (SD): 0.3μN) and were highly repeatable in a given microtissue as subsequent loading cycles were largely superimposable (SI 2).

Together stress relaxation and recovery responses indicated that microtissues modulate their tension in response to step changes in length towards their resting values. Importantly, however, microtissues did not completely relax upon a step strain, but rather seemed capable of maintaining residual stresses. Consequently, at each strain amplitude, microtissues reached a unique tension equilibrium that was independent of the loading history.

In order to further characterize the tension (T) relaxation and recovery responses, we searched for an appropriate model to describe these viscoelastic behaviors (fig. 2a and 2b, respectively). Classically engineers have modeled viscoelastic effects with a series of springs, described by their stiffness constants (k) and dashpots, described by their viscosity ^7^. In particular, the standard linear solid (SLS) model consists of a spring and a dashpot in series (ie. a Maxwell body) in parallel with another spring. Following a step change in length, this model predicts that tension will exponentially decay with a time constant, τ, towards a residual value (equation 2). The amplitude of this exponential function is given by the spring constant k_1_ while the residual stress is described by the spring constant k_2_. As a SLS model has been previously applied to early experiments on isolated cells^9–11^, 3D cell cultures^25^ and ex vivo tissue strips^7^, we begun by considering this model for describing microtissue mechanics. In that regard, figures 2a and 2b respectively show that a SLS model, could describe microtissue mechanics at short time-scales but failed to capture the long time tail in both relaxation and recovery responses.

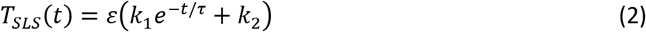

**Fig. 2:**
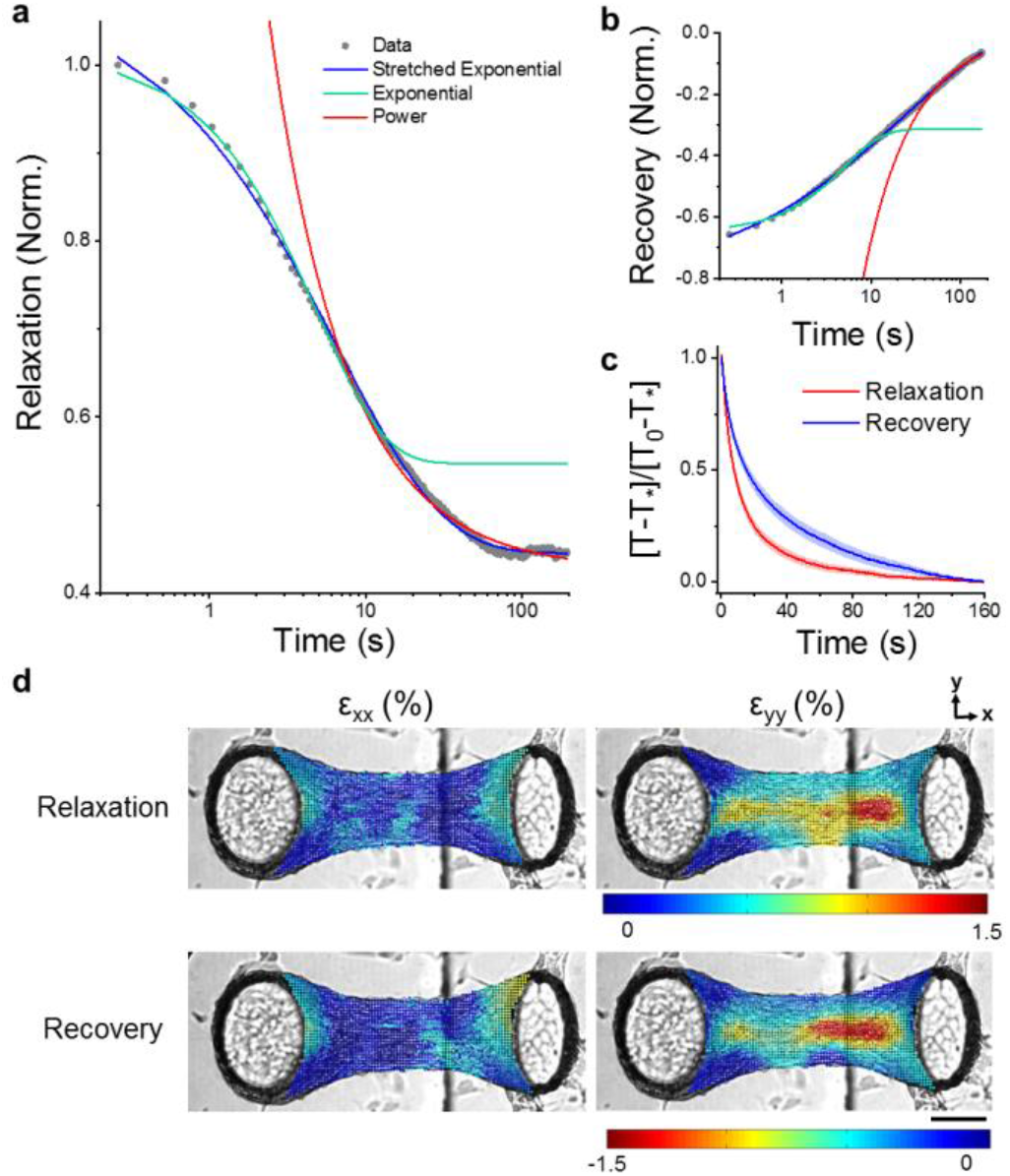
Stress relaxation and recovery in microtissues followed stretched exponential trajectories. Viscoelastic modeling of stress relaxation and recovery responses are respectively shown in (a) and (b). In both panals, representative data was normalized to the peak tension. Stretched exponentials capture relaxation and recovery behaviors over three decades of time. To assess the dynamics of the relaxation and recovery, responses were normalized to their amplitudes using tension measurements immediately following step length changes, T_0_, and after 160 seconds, T_*_. By normalizing responses in this manner, it is clear that relaxation occurred much quicker than stress recovery (c). Yet, stress relaxation and recovery appeared to share the same spatial locations within a given microtissue in terms of both remodeling in the longitudinal (ε_xx_) and transverse (ε_yy_) directions immediately following the change in length (d). The scale bar represents 100μm.

In contrast to an exponential model, slow relaxation and recovery at long timescales is exemplified by a power law rheological model. This second model predicts that stress relaxation/recovery should follow a power law with a dimensionless constant (β), which may describe a specific continuous distribution of discrete viscoelastic behaviors (equation 3). Since a long time tail behavior has been widely reported in single cells ^12–16^ and tissues ^6,18^, we next considered a power law model for describing microtissue mechanics. In that regard, a power law could capture the slow relaxation behavior of microtissues at long-timescales but predicted faster than experimental dynamics at shorter times

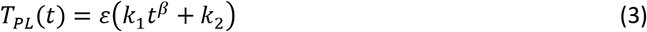

From considering these first two models, it was clear that in order to capture both the exponential stress decay at short-time scales and the slow relaxation at long time-scales, we needed a model that first appeared as an exponential but crossed over to a more slowly relaxing tail behavior at longer elapsed times. This behavior is characteristic of a so-called stretched exponential function (equation 4), which differs from a SLS model in that the time constant, τ, grows with a power law with elapsed time. In this model, one may interpret the power law constant, β, as a dimensionless descriptor of a specific distribution of discrete timescales, which broaden from a single timescale behavior as the value of β decreases from 1 towards 0. In other words, a stretched exponential function can appear by adding additional Maxwell bodies with a specific distribution of time constants to a SLS model. However, this method is an ineffective use of variables compared to a stretched exponential. A further discussion of stretch exponential model parameters and simulated stress relaxation responses are contained in SI 3.

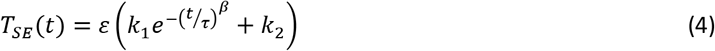

Although we have not come across previous reports that have modeled the viscoelastic behavior of cells or tissues with stretched exponentials, they have been widely used to describe relaxation in glassy, disordered systems that display a broad distribution of timescales ^35,36^. In regards to microtissues, however, a stretched exponential captured both stress relaxation and recovery data over three decades of time (R^2^>0.99) with average fitting constants summarized in table 1. Importantly, the creep that inherently accompanied our method of measuring microtissue tension relaxation was insufficient to significantly change either the model or the fitting constants (SI 4).

**Table 1:**
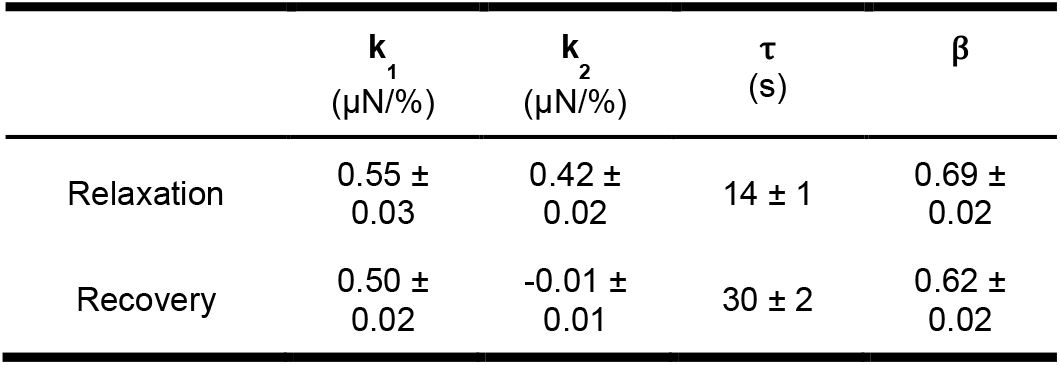
The stress relaxation and recovery fitting constants (N=110).

The amplitudes of the stretch exponential stress relaxation and recovery responses, given by k_1_ in table 1, agreed well each other (0.55 ± 0.03 vs. 0.50 ± 0.02 μN/%); P>0.05, repeated measures t-tests). Moreover, the spatial distributions of strain immediately following the step change in length, describing the locations of stress relaxation and recovery, were well correlated (fig. 2d). Because the regions that underwent stress relaxation were spatially and quantitatively linked to stress recovery, the viscoelastic response of microtissues likely occurred through a broad distribution of reversible remodeling events.

As for the spring constant k_2_ in our stretch exponential model, it characterizes the residual elastic stress that microtissues elastically store, as described above. On average microtissues had a residual elasticity of 0.42 ± 0.02 μN/%. The source of this elasticity is unclear but it significantly contributes to the mechanics of microtissues. It may arise from the contribution of the matrix, or perhaps indicates of the existence of a stress threshold up to which cytoskeletal elements can behave elastically but beyond which they yield. In recovery curves k_2_ is approximately equal to zero, indicating that microtissues fully recovered to their original prestress upon being returned to their initial length.

The dynamics of the relaxation and recovery responses are examined in figure 2c. In this figure, microtissue tension, T, has been normalized to the response amplitude measured across the initial 160 seconds. Normalizing the curves in this manner revealed that relaxation occurred much quicker than stress recovery (τ = 14 ± 1 vs. 30 ± 2 seconds, repeated t-test, P<0.001). Furthermore relaxation responses had a significantly, albeit slightly, higher power law constant (β = 0.69 ± 0.02 vs. 0.62 ± 0.02, repeated t-test, P<0.01). Therefore, although stress relaxation and recovery are spatially and quantitatively linked, the dynamics of these responses varied considerably; recovery occurred more slowly and over a slightly more broadly distributed set of timescales.

Taken together in this section we have described the viscoelasticity of microtissues and found that this behavior can be captured through a relatively efficient stretched exponential model. In the remainder of this article, we aimed to gain further insight into how the mechanics of microtissues are controlled by underlying molecular and structural mechanisms in the cells by first examining how this behavior is affected by strain, and secondly, how microtissue viscoelasticity changes under a range of pharmacological treatments targeting specific cytoskeletal elements.

### Microtissue viscoelasticity is nonlinear

To assess whether there were nonlinearities in the viscoelastic behavior of microtissues, stress relaxation and recovery measurements were performed at two different step strain amplitudes on the same microtissues. Shown in figure 3a are average response curves for both 3.3 ± 0.2 % and 10.9 ± 0.8% step strains (N=17). The changes to fitting constants for stress relaxation and recovery between these strain amplitudes are summarized in tables 2 and 3, respectively.

**Fig. 3:**
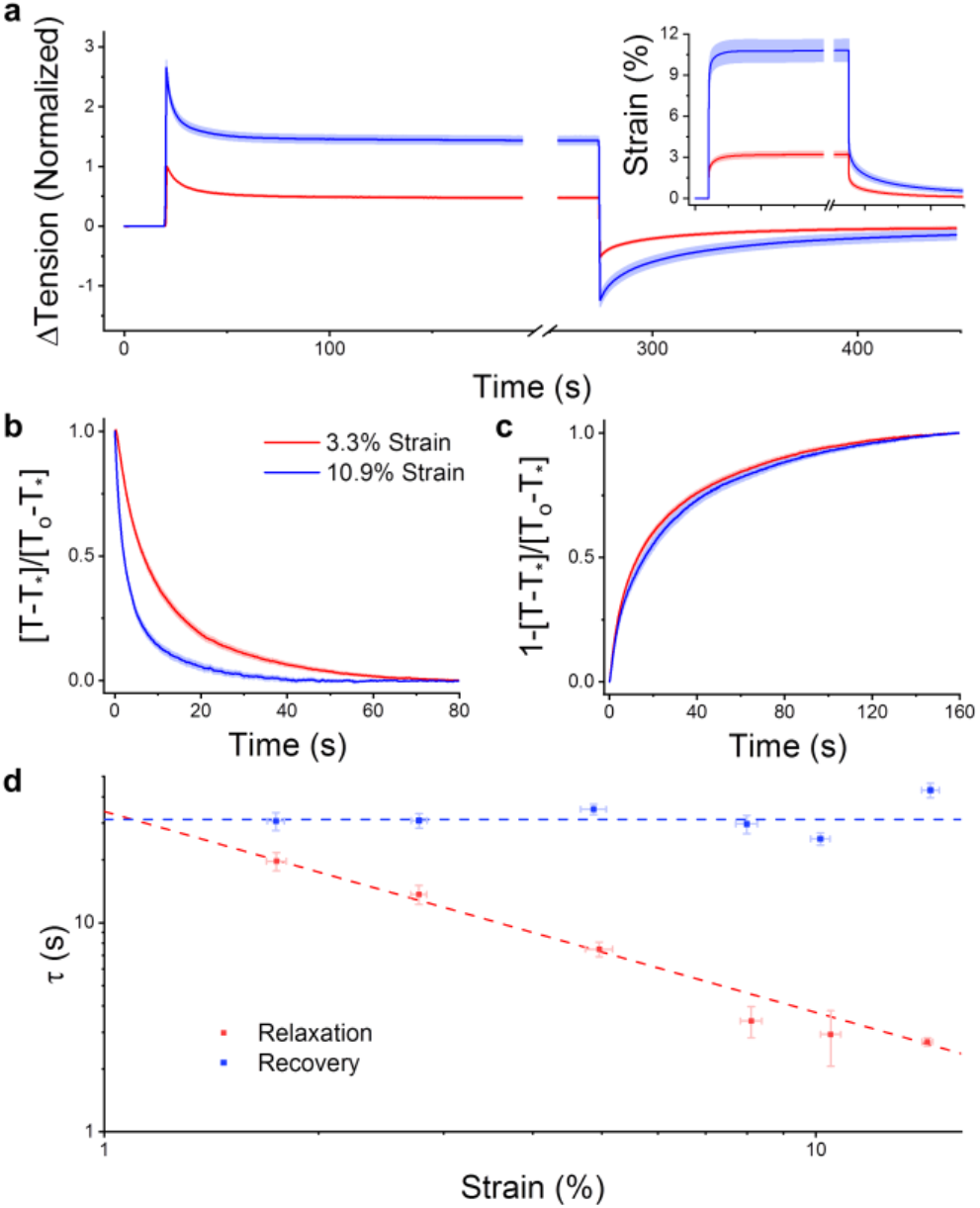
The rate of stress relaxation was nonlinear while recovery was strain invariant. To assess viscoelastic non-linearity, the step size was varied (insert). Stress relaxations for a 3.3 and 10.9% step are shown in (a) (N=17). After normalizing for response amplitude over the initial 80 seconds, (b) shows that the rate of stress relaxation increased with the strain amplitude. The rate of recovery was, however, invariant upon the step size (c). To further understand the nonlinearities in the viscoelastic behavior, 160 step strain experiments were completed across a range of strain amplitudes. Panel (d) shows that, as the strain amplitude increased, the time constant for stress relaxation decreased with an inverse power law with a near unity exponent, while in contrast, the rate of recovery was again invariant upon the step size.

**Table 2:**
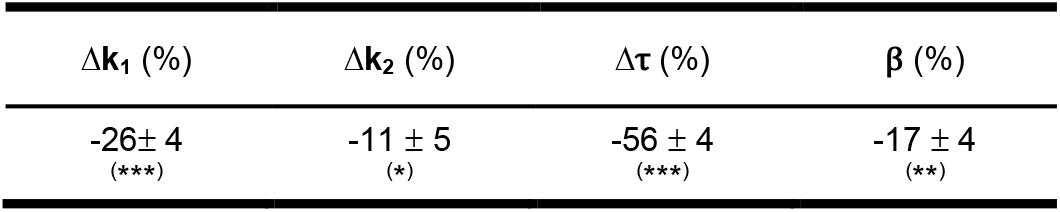
The changes to stress relaxation fitting constants measured at 10.9% vs. 3.3% strain. Values were compared with repeated measures t-tests (N=17) (* P<0.05, ** P<0.01, *** P<0.001).

**Table 3:** The changes to stress recovery fitting constants measured at 10.9% vs. 3.3% strain. Values were compared with repeated measures t-tests (N=17) (** P<0.01).

| $\Delta k_1$ (%) | $\Delta \tau$ (%) | $\beta$ (%) | $\Delta T_{f,i}$ ($\mu\text{N}$) |
| --- | --- | --- | --- |
| $-28 \pm 5$<br>(**) | $15 \pm 7$ | $-2 \pm 3$ | $-0.4 \pm 0.2$ |

A larger step strain amplitude decreased both k_1_ and k_2_ spring constants for relaxation responses respectively by 26 ± 4 and 11 ± 5 % (repeated measures T-tests; for k_1_: P<0.001, for k_2_: P<0.05). Likewise, the k_1_ spring constant fitted to recovery responses decreased by 28 ± 5% (repeated measures T-test; P<0.01). These responses were indicative of a strain amplitude softening behavior that followed from abrupt loading.

The relaxation responses were normalized in figure 3b to show how the dynamics varied with the strain amplitude. In that regard, a larger step strain decreased τ by 56 ± 4% and β by 17 ± 4% (repeated t-tests; for τ: P<0.001, for β: P<0.01). Therefore, by increasing the step size, microtissues relaxed more quickly and slightly over a more broadly distributed set of timescales.

In contrast to stress relaxation responses, the time and power law constants of stress recovery were step strain amplitude invariant (fig. 3c) (repeated t-tests; both P>0.05). Moreover, they did not depend upon the strain at which the microtissue recovered (SI 5). Therefore, whereas the rate of stress relaxation in microtissues was nonlinear with strain amplitude, stress recovery was a linear process.

To further define the relationships between stretch exponential fitting constants and the strain amplitude, 160 responses with step strains ranging from 1.75 ± 0.06% to 14.3 ± 0.2% were discretized into strain bins and analyzed with regression models. In accordance with repeated measures observations, figure 3d shows that the relaxation time constant decreased with the strain amplitude, and surprisingly followed a power law dependence (non-linear regression: R^2^=0.96 P<0.001). Further in agreement, the recovery time constant was strain-invariant (linear regression: P>0.05). The remaining fitting constants to relaxation and recovery responses are respectively summarized in SI 6 and 7.

Thus far in this section, we have shown that microtissue mechanics varied considerably with the imposed strain amplitude. To link these mechanical behaviors to changes in the spatial remodeling response, the strain distribution was characterized over time immediately starting after the change in length. Figure 4a and b respectively show the integrated raw and normalized remodeling in the transverse direction in representative microtissues over 100 seconds of stress relaxation and recovery. As with the fitting constants, the spatial distribution of microtissue remodeling following a step change in length varied considerably with strain. In that regard, remodeling during stress relaxation and recovery in the transverse direction increased with step size and became transversely focused to the center of the tissue but longitudinally diffuse, spanning the entire tissue length.

**Fig. 4:**
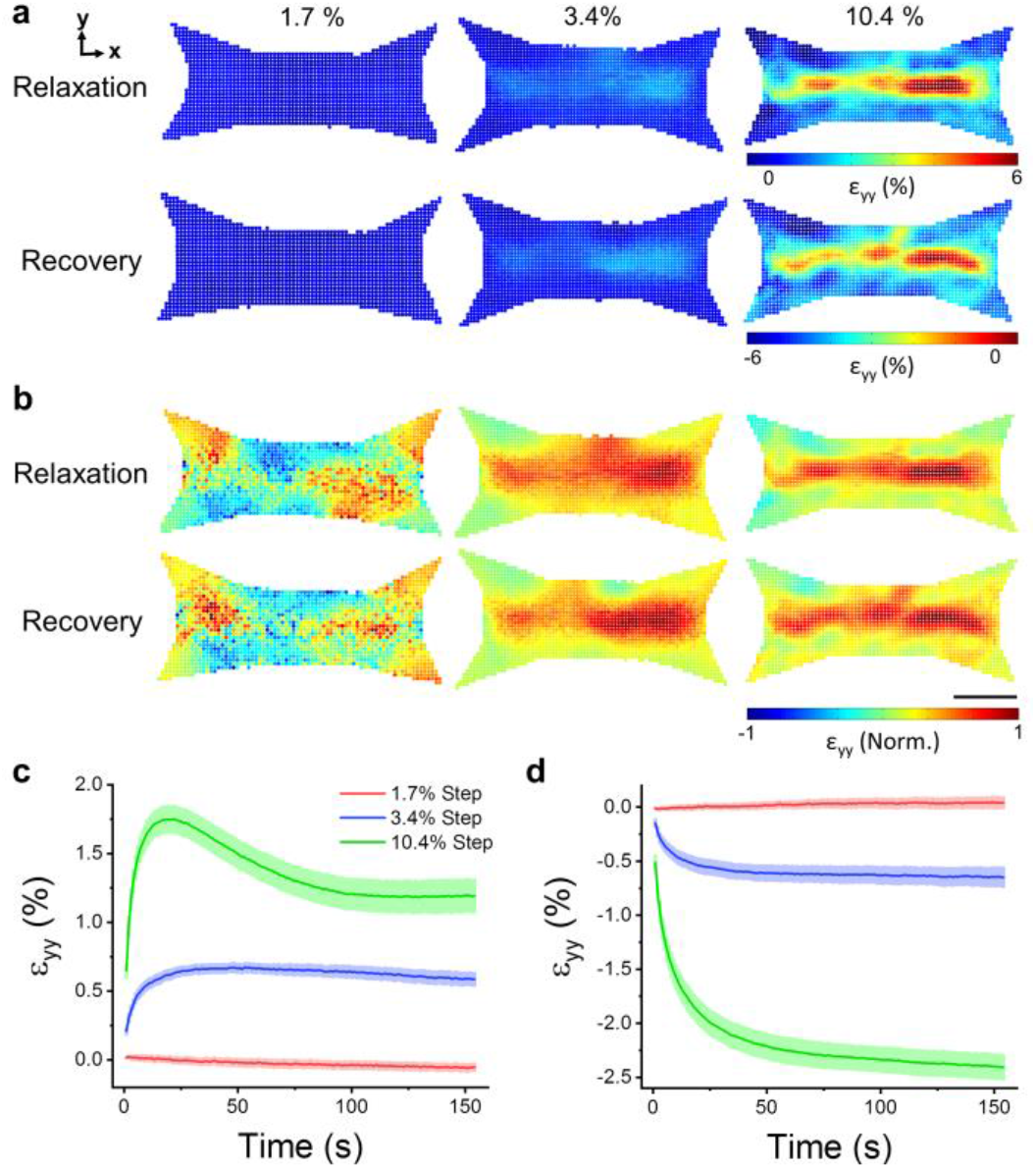
The locations of remodeling following changes to microtissue length were strain-dependent. Spatial distributions of relaxation and recovery in the y-direction immediately after various step sizes are shown in (a). The distributions are normalized to three standard deviations outside of the absolute mean value in (b). The scale bar represents 100μm. The average (N=6) strain in the transverse direction that occurred during stress relaxation and recovery is plotted against elapsed time in (c) and (d), respectively.

To compute a mean metric of the integrated relaxation response through time, transverse strain fields were averaged across the region of interest (fig. 4c). Following a small step strain (1.7 ± 0.2%), microtissues showed comparably little relaxation in the transverse direction. In comparison with an intermediate step strain (3.4 ± 0.2%), microtissues relaxed to reach a new equilibrium. Lastly, with a large step strain (10.4 ± 0.5%), microtissues quickly relaxed, then contracted to a smaller degree, and eventually reached an equilibrium. This intermediary contraction response to a large step strain was not seen in tension measurements and in longitudinal remodeling (SI 8).

Compared to relaxation responses (fig. 4c), mean recovery remodeling showed slower dynamics as expected from mechanical measurements (fig. 4d). Yet, relaxation and recovery following small and intermediate step strains shared similar spatial distributions and amplitudes with opposing signs, indicating that the gross structural remodeling that occurred after a step strain was a reversible process. In contrast however, transverse recovery remodeling following a large step strain was comparably greater than the final equilibrium in relaxation curves because of the delayed contraction.

In comparison with the remodeling in the transverse direction, remodeling in the longitudinal direction was less extensive being constrained by the cantilevers. Longitudinal remodeling was also highly tissue-specific and mainly located near the fronts of the cantilevers rather than the body of the tissue. Since final relaxation and recovery longitudinal strain fields shared similar amplitudes and microtissue tension fully recovered, this behavior likely did not indicate tissue detachment from the cantilevers (SI 8). Rather the longitudinal remodeling response more likely reflected stress concentrations formed around the front of the cantilevers created through a reduced fraction of longitudinally oriented actin filaments (fig. 1b)

### Pharmacological behaviors

To assess the role of several key cellular proteins in governing microtissue viscoelasticity, stress relaxation and recovery were measured following various pharmacological treatments meant to either disrupt the cytoskeleton or alter actomyosin activity. Normalized responses and fitting constants are shown in fig. 5a and table 4, respectively. Each fitting constant was normalized and compared with paired t-tests to their own pre-treatment control. DMSO was used as a loading control and did not affect either relaxation or recovery curves. Furthermore tension measurements taken prior to and following step strain experiments did not vary with any treatment, indicating that stress relaxations were fully recoverable under each condition and that incubation periods were sufficient to reach an equilibrium state.

**Fig. 5:**
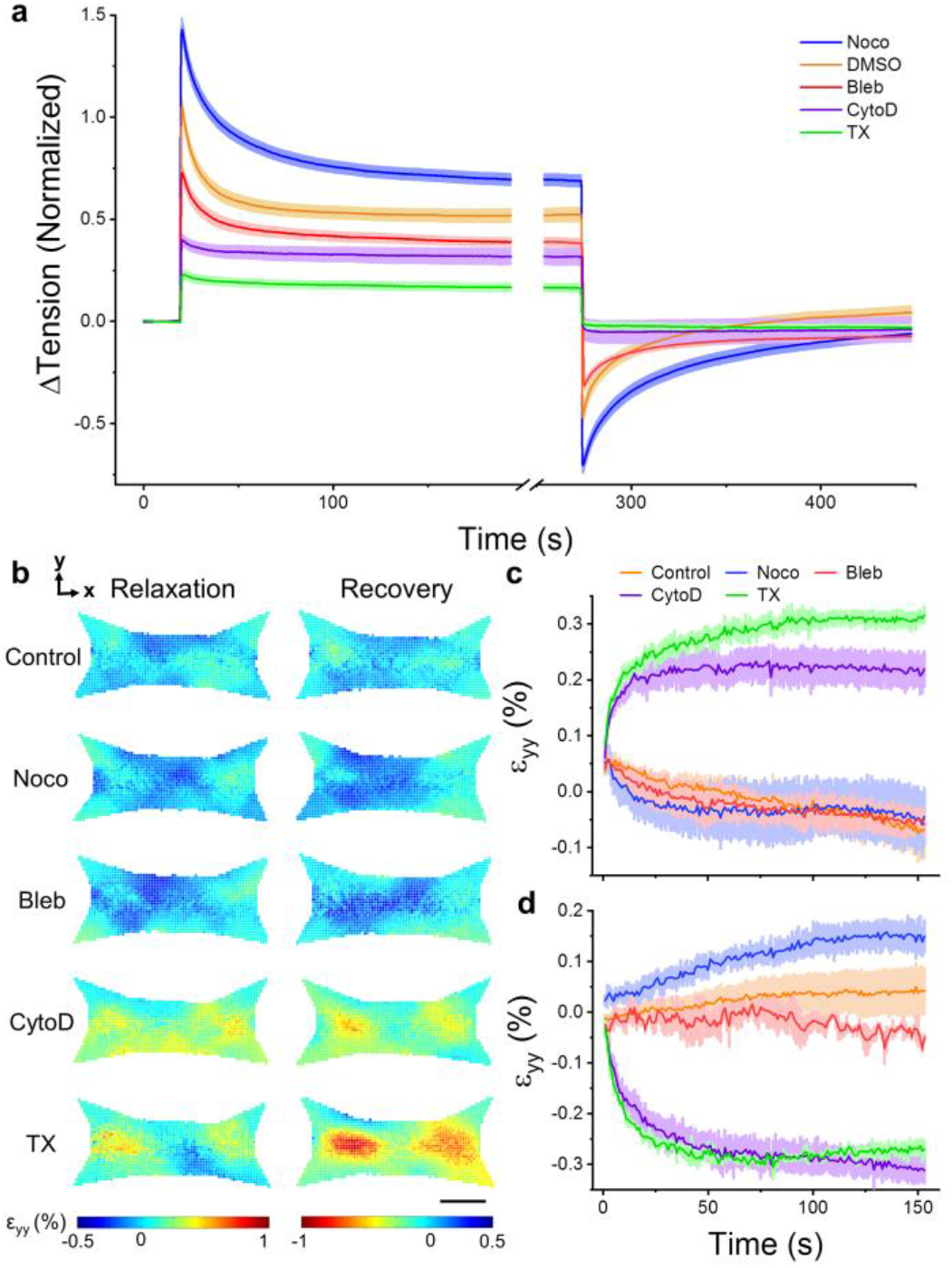
Stress relaxation and recovery varied with pharmacological disruption of the cytoskeleton and myosin inhibition. Depolymerization of microtubules (Noco) increased stress relaxation and recovery responses, whereas myosin inhibition (Bleb), actin depolymerization (CytoD) and decellularization (TX) decreased relaxation and recovery (a). DMSO, a loading control, had no difference compared to no treatment. Microtissues only ever received a single treatment and each treatment was normalized to its pretreatment control. The spatial distributions of relaxation and recovery in the transverse direction following a small (1.7%) change in length are shown in (b). The scale bar represents 100μm. The average (N=3) strains in the transverse direction during stress relaxation and recovery are shown in (c) and (d), respectively.

**Table 4:** The pharmacologically induced changes to stress relaxation and recovery fitting constants. Each treatment was compared to its pretreatment control using paired t-tests (* P<0.05, ** P<0.01, ***P<0.001).

| | $\Delta T_{\text{base}}$<br>($\mu\text{N}$) | Relaxation | | | | Recovery | | | |
| --- | --- | --- | --- | --- | --- | --- | --- | --- | --- |
| | | $\Delta k_1$<br>(%) | $\Delta k_2$<br>(%) | $\Delta \tau$<br>(%) | $\Delta \beta$<br>(%) | $\Delta k_1$<br>(%) | $\Delta \tau$<br>(%) | $\Delta \beta$<br>(%) | $\Delta T_{f,i}$<br>($\mu\text{N}$) |
| Noco<br>(N=24) | $3.7 \pm 0.4$<br>(***) | $40 \pm 9$<br>(***) | $37 \pm 9$<br>(***) | $10 \pm 12$ | $11 \pm 5$ | $96 \pm 14$<br>(***) | $87 \pm 25$<br>(**) | $-7 \pm 4$<br>(*) | $0.03 \pm 0.07$ |
| DMSO<br>(N=21) | $-0.1 \pm 0.2$ | $6 \pm 6$ | $8 \pm 7$ | $4 \pm 9$ | $8 \pm 8$ | $10 \pm 9$ | $5 \pm 13$ | $8 \pm 5$ | $0.17 \pm 0.09$ |
| Bleb<br>(N=16) | $-2.8 \pm 0.5$<br>(***) | $-27 \pm 7$<br>(**) | $-15 \pm 5$<br>(*) | $14 \pm 10$ | $-2 \pm 9$ | $-32 \pm 5$<br>(***) | $-27 \pm 8$<br>(**) | $7 \pm 4$ | $-0.1 \pm 0.06$ |
| CytoD<br>(N=8) | $-5.0 \pm 0.4$<br>(***) | $-74 \pm 10$<br>(***) | $-47 \pm 12$<br>(***) | - | - | $-112 \pm 6$<br>(***) | - | - | $-0.1 \pm 0.1$ |
| TX<br>(N=7) | $-8.8 \pm 0.8$<br>(***) | $-86 \pm 6$<br>(***) | $-65 \pm 7$<br>(**) | - | - | $-109 \pm 10$<br>(***) | - | - | $-0.1 \pm 0.2$ |

Firstly, consistent with our understanding that microtubules act as compressive struts that oppose actomyosin activity ^37^, microtubule depolymerization with nocodazole increased the resting tension in microtissues by 3.7 ± 0.4μN (P<0.001), as well as spring constants for stress relaxation (k_1_ and k_2_ both P<0.001) and recovery (k_1_ P<0.001). While the time or power law constants for stress relaxation (τ and β both P>0.05) were not affected, microtubule depolymerization increased the stress recovery time constant by 87 ± 25 % and decreased the power law constant by −7 ± 4 % (P<0.01 and P<0.05, respectively). This suggests that although the relaxation rate is not affected, the recovery rate and, more modestly, the broadness of the distribution of timescales in the response may be altered through the regulatory action of microtubules on acto-myosin activity.

Secondly, as expected, myosin inhibition with blebbistatin decreased resting microtissue tension by −2.8 ± 0.5 μN (P<0.001), and reduced relaxation and recovery spring constants (various P values). Similar to microtubule depolymerization, blebbistatin treatment did not affect either the relaxation time or power law constants (τ and β P>0.05). Blebbistatin treatment did decrease the recovery time constant by −27 ± 8% (P<0.01), which indicates that myosin motor activity, also in part, determines the recovery speed. However, unlike microtubule depolymerization, myosin inhibition did not affect the power law constant (P<0.0), and thus does not determine the broadness of the distribution of timescales.

Lastly, actin depolymerization (cytochalasin D) and decellularization (triton-X) had similar effects on the mechanical behavior of microtissues. Both treatments decreased resting microtissue tension (−5.0 ± 0.4μN and −8.8 ± 0.8μN, respectively; both P<0.001), as well as relaxation and recovery spring constants (various P values). In fact neither treatment displayed observable stress relaxation or recovery responses, and for this reason, it was not possible to accurately fit time and power law constants. Nevertheless, these responses revealed that remodeling in actin microfilaments is chiefly reasonable for the viscoelastic behavior of microtissues.

To examine how microtubules, actin microfilaments, myosin motor activity and the extracellular matrix contributed to microtissue remodeling during relaxation and recovery, strain fields were assessed following a small step strain (1.7 ± 0.2%) with the pharmacological treatments. Microtubule depolymerization and myosin inhibition behaved similarly to control microtissues in their transverse (fig. 5b-d) and longitudinal (SI 9) remodeling responses. In contrast, actin depolymerization and decellularization induced remodeling responses that were more consistent with a larger step strain in terms of the spatial distribution, amplitude and dynamics of the strain fields, apart from lacking an intermediary contraction response. This finding seems to implicate the depolymerization of actin and the contribution from the matrix in the gross remodeling behavior and nonlinear dynamics observed with large strains.

## Discussion

In this article, we assessed the dynamic mechanical behavior of physiologically relevant 3D microtissue cell cultures. As expected in living active matter, microtissues displayed timescale-varying mechanics. In response to a step increase in length, the tension in microtissue cultures quickly rose and then relaxed to a new equilibrium stress, resembling a viscoelastic solid. Then when microtissues were returned to their initial lengths by a recovery strain, their tension quickly dropped below initial measurements, and then slowly, yet fully, recovered. As these responses differed from the widely observed weak power laws that describe the relaxation behavior of cells in 2D culture ^12–16^, our findings in 3D microtissues may reflect additional considerations necessary to completely understanding the viscoelastic response of living tissues. In that regard, while our field has largely been focused on characterizing the mechanical response of individual cells, tissues mechanics is perhaps instead generated by an aggregate of active cytoskeletal dynamics in a heterogeneous population of cells, and through interacting with a soft largely elastic 3D matrix.

In particular, unlike previous work in 2D cell culture, we found that tensile stress relaxation and recovery in microtissues followed stretched exponential functions. To the best of our knowledge, stretched exponentials have not been previously used to describe the viscoelastic behavior of cells or cell cultures. Still, however, our findings are in close agreement with previous reports from bulk fibroblast populated collagen gels ^25,27^. In that regard, we both reported characteristic time constants of similar magnitudes with time-dependencies that are more broadly distributed than what can be explained by a single exponential function (ie. a SLS model).

On the contrary, there does exist a strong precedent for the use of stretched exponentials to describe relaxation in disordered condensed matter physics. In fact both polymer networks ^38^ and glasses near their transition temperature ^35,36^ relax according to stretch exponentials. Notably, such glassy polymer response has already been linked to the mechanical behavior of the cytoskeleton ^39^. Furthermore, our average values of β (0.64 ± 0.01 and 0.61 ± 0.01 for relaxation and recovery, respectively) are surprisingly close to an universal experimental value of 0.6 in glasses dominated by Brownian motion ^40^, which is theoretical justified by a so-called ‘trapping model’ ^41^. Mathematically this model is beyond the scope of this work but it is relatively easy to conceptualize. In a trapping model, the material consists of randomly distributed traps. During relaxation, these traps capture molecules that diffuse through the material, and as the traps are filled, the remaining molecules must travel longer to find unoccupied traps. Thereby, the relaxation rate falls with a power law given by the dimensionality of the model: in 3D environments, β=0.6.

Although a trapping model can be a great tool to conceptualize stretched exponentials, the question remains whether we can draw out any physiological meaning or link any mechanism(s) to account for the stretched exponential relaxation and recovery in microtissue behavior. Firstly, similar to power laws, stretched exponentials can be expanded into a superposition of exponential functions with a nontrivial distribution of relaxation times ^42^. In this respect, the power law constant describes the broadness of the distribution. It should not be overly surprising that relaxation processes exist across a broad spectrum of timescales in something as complex and heterogeneous as the network of cytoskeletal proteins inside single cells, and in a more collective sense, when considering an aggregate of cells and extracellular matrix. Thus, our fitting constants may be interpreted as a general description of microtissue behavior reflecting the broad viscoelastic heterogeneity in the cytoskeleton and amongst cells rather than capturing a specific process. What may be more surprising was that the same model, describing an inter-related set of exponential functions, could be applied over a range of pharmacological treatments and strain amplitudes with only small changes to the power law constant, while changes to the other fitting constants were large. At the moment, an explanation for this behavior remains unclear. Perhaps future work in generating a more mechanistic modeling approach may shed further insight into the stretch exponential behavior of microtissues and their power law constant.

In addition to displaying stretch exponential dynamics, we found that the relaxation spring constants (k_1_ and k_2_) of microtissues remained tightly coupled throughout an assortment of pharmacological treatments targeting specific proteins, and as well in measurements at different step strain amplitudes. In that regard, all treatments followed a single linear relationship (fig. 6a). Although assigning specific meaning to these spring constants would be speculative, our results indicated they both largely depended upon the actin cytoskeleton, and to a lesser extent myosin activity, the presence of microtubules, and the elasticity of the matrix. Previously this coupling behavior between energy loss and residual elasticity in tissues has been interpreted as evidence suggesting that residual elastic and dissipative stresses are borne from the same origin(s) ^43,44^. Even so, it is surprising that targeting different cytoskeletal elements can produce responses that follow a single universal relationship, unless an understanding of mechanisms at a hierarchy beyond the single protein-level is required. In other words, rather than following responses in single molecules, as our field has been studying in a traditional reductionist approach, the viscoelasticity of cells and tissues is perhaps more influenced by the association and interaction of proteins with each other ^45^. For example, through bundling collagen filaments, cells may control the residual elasticity of the matrix, and thereby, any change to acto-myosin dynamics will also be seen as a change in the stiffness of the matrix. Further evidence in support of this view has also been reported through various universalities in single cell mechanics ^13,37,46,47^.

**Fig. 6:**
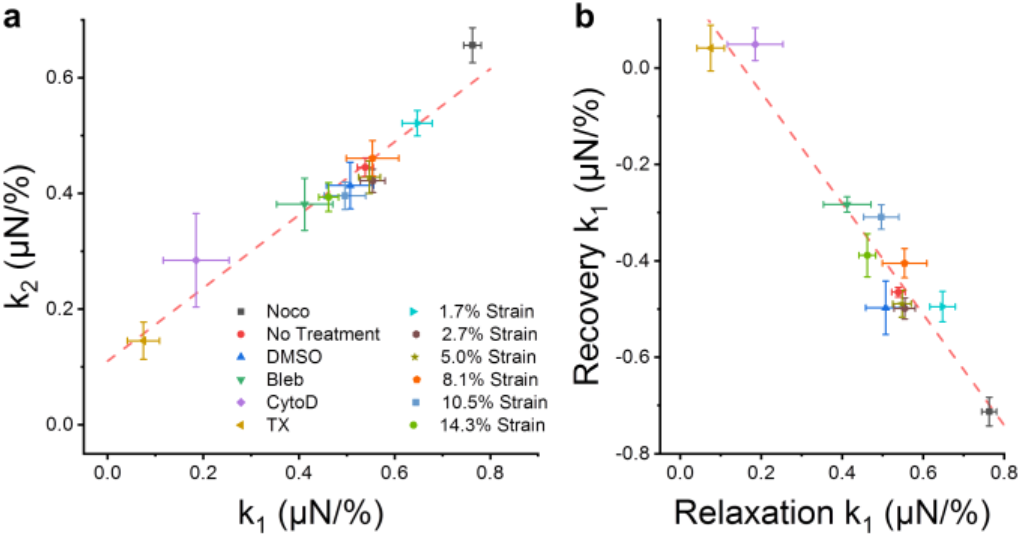
Underlying relationships in microtissue mechanics. The k_1_ and k_2_ stress relaxation spring constants were linearly coupled throughout an assortment of pharmacological treatments and step lengths (a) (linear regression; P<0.001 R^2^=0.94). Likewise the amplitude of stress relaxation was coupled to the amplitude of stress recovery (b) (linear regression; P<0.001 R^2^=0.92).

The degree of stress relaxation was also strongly coupled to the stress regained during recovery (fig. 6b). Furthermore, for the most part, microtissue remodeling during relaxation and recovery occurred with similar spatial distributions and comparable amplitudes. Together these results suggest that stress relaxation and recovery depended upon reversible remodeling in the same elements within the microtissue. That being said, a notable exception was observed in the transverse remodeling response to a large amplitude (10%) strain, where a delayed contraction followed stress relaxation. This contractile behavior following a large step strain is consistent with stretch activated contraction responses previously reported in isolated fibroblast cells ^48^ and 3D cultures ^49^, and it seems to act separate from stress relaxation and recovery, causing permanent remodeling.

In contrast to the amplitudes, the dynamics of the relaxation and recovery responses differed greatly. Relaxation rates were nonlinear with step strain amplitude, decreasing with what appeared to be a power law, but were relatively invariant with microtubule depolymerization or myosin inhibition. In contrast, recovery rates were invariant with step strain amplitude but increased with microtubule depolymerization and decreased with myosin inhibition. This suggests that myosin contributed only as a passive cross-linker during relaxation, while active myosin crossbridge cycling was partially responsible for stress recovery. Still however, the actin cytoskeleton was the principal source of the viscoelastic behavior of microtissues since depolymerization of actin filaments had the biggest effect on step strain responses outside of decellularization. There also appeared to be some passive component to the recovery as there was a small remodeling response upon returning samples with actin-depolymerized cells and as well as in decellularized microtissues to their initial lengths. Interestingly, this remodeling was consistent with larger step strains, suggesting that the loss of the actin cytoskeleton and how the cells associate with the matrix may be in part be responsible for the non-linear relaxation rates.

It is not uncommon for relaxation in soft biological tissues to be nonlinear. For example, nonlinearities have been previously reported in lung^50^, heart valve^51^, ligament^52^, and muscle^53^. Recently, nonlinear relaxation rates in cells were argued to arise from a poroelastic effect occurring through the redistribution of the viscous cytosolic fluid between local regions within the network of cytoskeletal filaments^17^. It is, however, debatable that the same poroelasticiy effect is the main determinant in our measurements, as our time-constants differed more than a decade from those previously reported. Moreover, poroelasticity does not offer an explanation as to why the recovery rates were linear with strain amplitude and much slower than relaxation rates.

An alternative mechanism conceives stress relaxation in cells as a friction force produced by cytoskeletal filaments sliding smoothly past one another ^54^. The main support for this hypothesis is that local shear measurements on cells in 2D culture follow the structural damping law; the ratio of dissipative to elastic stresses is invariant of the time-scale at which they are assessed ^12,13^. This hypothesis, however, does not offer any explanation as to the nonlinearity observed in relaxation rates and the imbalance with recovery dynamics. Furthermore it is not compatible with either standard linear solid viscoelasticity or a stretched exponential, as both models produce characteristic rate-dependencies that contradict structural damping. Interestingly, in contrast with local shear measurements, characteristic rate-dependencies similar to our observations in microtissues have been previously reported in individual cells^11^ and 3D cultures^25^ when probed in tension at a length-scale of the entire cell/culture. It would still be of interest, however, in future work to show that the frequency response of microtissues is consistent with their stretched exponential relaxation in the time domain.

A last possible mechanism to explain the viscoelastic response in microtissues is that stress relaxation and recovery reflects the rupturing and reforming of bonds within the cytoskeleton (i.e. within actin filaments and between actin and its crosslinkers), and between the cells and the extracellular matrix. In support of this hypothesis, there is a strong precedent that mechanical stretch depolymerizes actin filaments^46,55–57^ and perturbs myosin binding^58–62^. Indeed, we have shown here that microtissues strain-soften in an amplitude dependent manner. Further in regard to this hypothesis, the imbalance in microtissue relaxation and recovery rates can simply be explained by differences in bond destruction and formation rates; it is not unreasonable to suspect that it takes less time to pull bonds apart than for proteins in a cell to diffuse through Brownian motion, correctly reorient themselves, and then lastly reform chemical bonds, especially if additional enzymes (ie. profilin) and molecules (ie. ATP) are needed. Furthermore, nonlinearities in relaxation behavior have previously been captured by models in which stress is relieved through sequential rupturing of Maxwell bodies, where each micro-yield event passes the stress on to other regions of the tissue ^63,64^. Results from these models give an exponential with a long-tail power law, however it is possible that chemical bonds have a finite yield strength, and therefore in actuality curves may flatten, behaving as stretched exponentials. Admittedly, however, this model has yet been used explain nonlinearities beyond quasi-linearity, such as the inverse power law between relaxation rate and strain amplitude that we observed.

## Conclusions

The viscoelastic behavior of microtissues likely does not come down to a single physiological mechanism, but rather it is an amalgamation of many physical remodeling events occurring at inter-related time and length scales. Accordingly, the measured stress relaxation and recovery of microtissues followed generalized stretch exponential behavior, differing from the power law rheology seen in isolated cells. We further found that stress relaxation rates were nonlinear and largely imbalanced with a linear recovery response. With that said, the contributions of specific cytoskeletal elements (actin, myosin, microtubules) to microtissue mechanics did not qualitatively differ from our previous understanding of their roles in the mechanical behavior of cells grown on 2D substrates; actin was predominantly responsible for the viscoelasticity of microtissues. Taken together, however, our results collapsed onto a single relationship between residual elastic stresses and relaxation amplitudes, indicating that the mechanics of microtissues may follow an universal behavior set not by the role of individual proteins but rather by the physical architecture of cells and their surrounding matrix as a whole. To conclude, the assessment of dynamic mechanics of microtissues has yielded further insights into how tissues gain their mechanical properties from cells, and from the distribution and interaction of their cytoskeletal proteins. This knowledge is critical to a full understanding of how physical forces are sensed and regulate cell behavior in health and disease.

## Additional Information

### Supplementary Information

See supplementary information for methodological validations and further step strain experimental results.

## Acknowledgements

M.W. is supported by OGS (Ontario Graduate Scholarship). The authors acknowledge support from individual NSERC Discovery Grants (M.G. and A.E.P.). M.G. and A.E.P. acknowledge the Canadian Foundation for Innovation.

## Data Availability

The data generated during the current study is available from the corresponding author upon reasonable request.

