## Supplemental Data for "Time dependent stress relaxation and recovery in mechanically strained 3D microtissues"

*Condensed Running Title: microtissue viscoelasticity*

Matthew Walker[1], Michel Godin[2,3,4], James L. Harden[2,5], Andrew E. Pelling[1,2,6,7]*

[1]Department of Biology, Gendron Hall, 30 Marie Curie, University of Ottawa, Ottawa, ON, K1N5N5 Canada
[2]Department of Physics, 150 Louis Pasteur pvt., STEM Complex, University of Ottawa, Ottawa, ON K1N 6N5 Canada
[3]Department of Mechanical Engineering, Colonel By Hall, 161 Louis Pasteur, University of Ottawa, Ottawa, ON K1N6N5 Canada
[4]Ottawa-Carleton Institute for Biomedical Engineering, Colonel By Hall, 161 Louis Pasteur, University of Ottawa, Ottawa, ON K1N6N5 Canada
[5]Ottawa Institute of Systems Biology, University of Ottawa, Ontario K1H 8M5, Canada
[6]Institute for Science Society and Policy, Simard Hall, 60 University, University of Ottawa, Ottawa, ON, K1N5N5 Canada
[7]SymbioticA, School of Human Sciences, University of Western Australia, Perth, WA, 6009 Australia

*Keywords:*
Microtissue, Viscoelasticity, Cell mechanics, 3D cell culture, Microfabrication

* Author for correspondence
Andrew E. Pelling
150 Louis Pasteur pvt.
University of Ottawa
Ottawa, ON K1N 6N5
Canada

Web: http://www.pellinglab.net

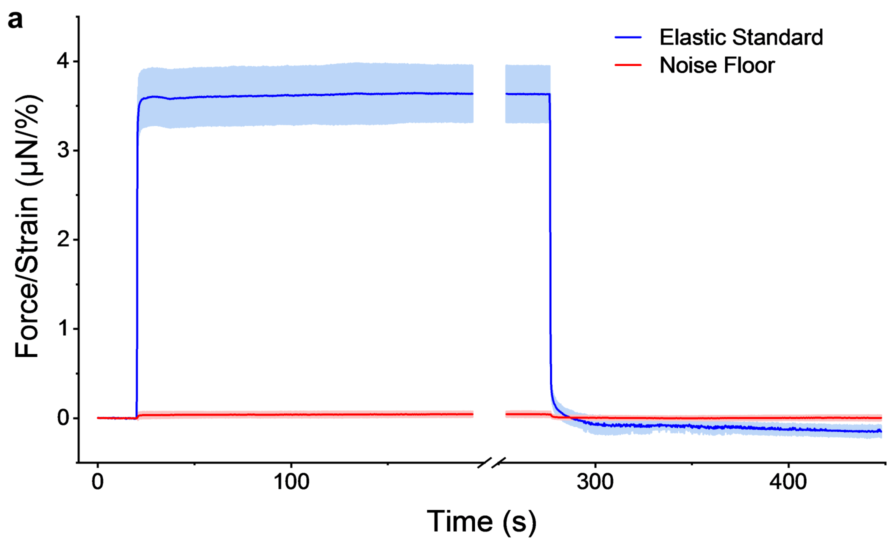


**SI 1: MVAS-force validation for step strain measurements.** Step strain experiments were performed without an attached load (N=5), to measure the noise floor, and with a polymerized 70x15µm strip of PDMS (N=3), to act as an elastic standard (a). The noise floor was much smaller than microtissue force measurements. Furthermore, as expected, there was no relaxation or recovery response with measurements of the elastic standard or the noise floor. The errors represent the standard deviation.

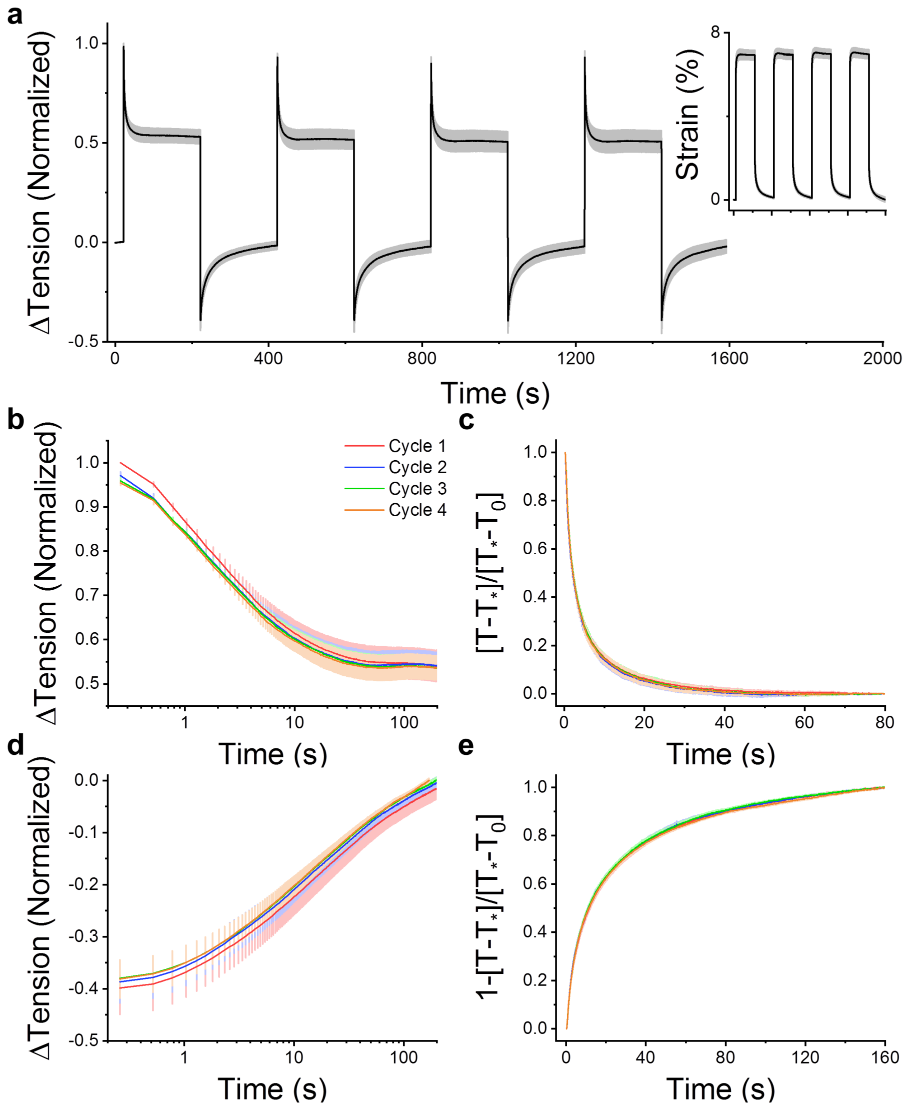


**SI 2: Microtissue viscoelastic behavior is highly repeatable.** Four step strain experiments were completed sequentially on the same microtissues (N=6) (a). Stress relaxations shared similar amplitudes (b) and rates (c) and so did stress recoveries (d and e). There were no significant differences to any of the fitting constants ($k_1$, $k_2$, τ, or β) for relaxation or recovery (repeated measures 1-way ANOVA; $P>0.05$).

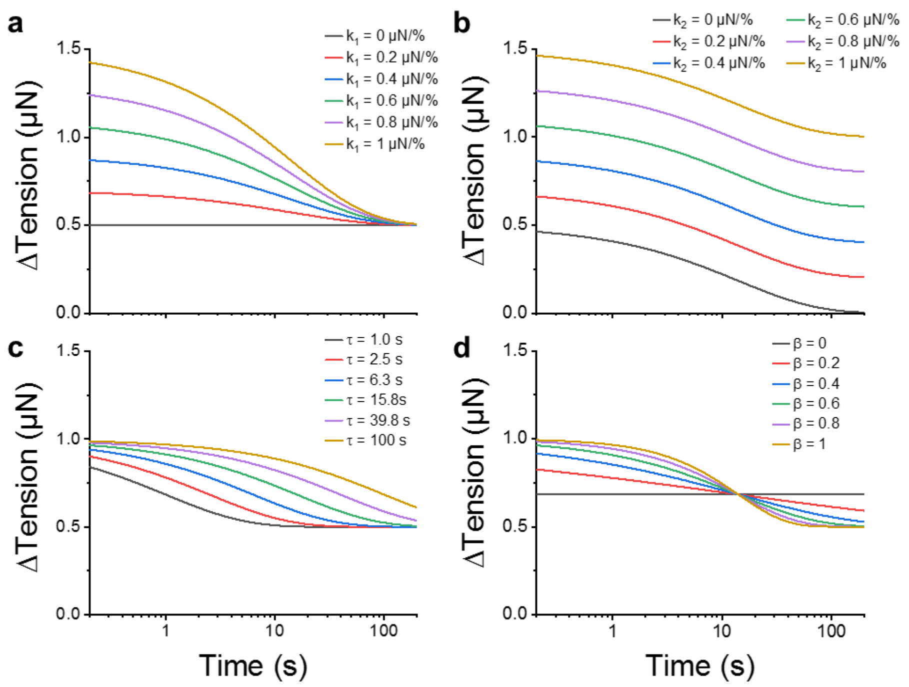


**SI 3: Simulated stretch exponential relaxation responses.** Each panel in this figure was simulated by varying one model parameter: $k_1$ (a), $k_2$ (b), $\tau$ (c) and $\beta$ (d). The $k_1$ and $k_2$ spring constants respectively describe the amplitude of the relaxation response (a) and the residual stress (b). $\tau$ is the time constant and determines the dynamics of the response (c). For a stretched exponential, the time constant grows with a power law constant, $\beta$, which is a dimensionless constant that captures a specific distribution of timescales (d). When $\beta$=1, the model behaves as a SLS with a single time constant. Whereas when $\beta$ approaches 0, the distribution of timescales broadens.

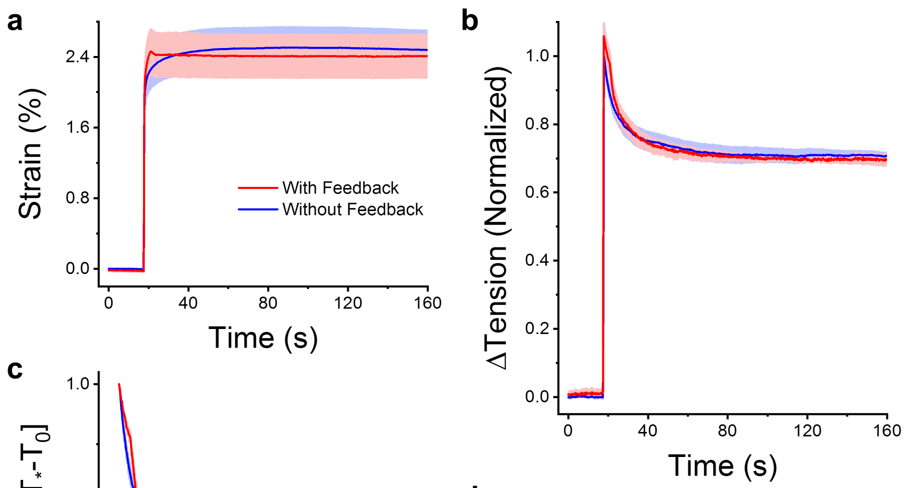


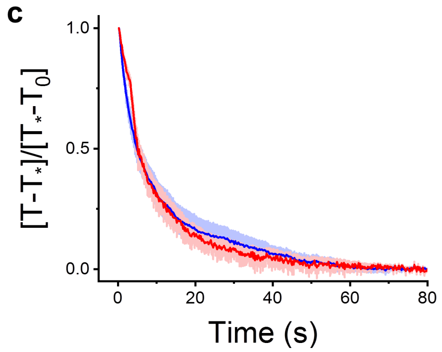


**d**

| | **$k_1$** (norm.) | **$k_2$** (norm.) | $\tau$ (s) | $\beta$ |
|---|---|---|---|---|
| With feedback | 0.26 ± 0.06 | 0.39 ± 0.05 | 8 ± 0.8 | 0.7 ± 0.1 |
| Without feedback | 0.20± 0.02 | 0.39 ± 0.04 | 8 ± 2 | 0.7 ± 0.1 |

**SI 4: Microtissue creep does not affect assessment of stress relaxation behavior.** Because our method of measuring tissue tension through cantilever deflection inherently causes some creep in the tissue length, we assessed how our measurements were affected by first doing a step strain experiment as normal (without feedback) and then by redoing the step strain experiment on the same microtissues but while modulating the vacuum pressure so to get rid of the creep response in the original experiment (with feedback) (a) (N=4). The changes to tension (b) and the rate of relaxation (c) were comparable without and with modulating the vacuum pressure to remove creep. Furthermore both relaxation responses followed stretched exponential trajectories; the fitting constants are in (d). As expected, the $k_1$ constant was slightly higher, but not significantly ($P>0.05$, repeated measures t-test), when there was no creep in microtissue length. There was no change in $k_2$, the time constant, or the power law constant (repeated measures t-tests; $P>0.05$).

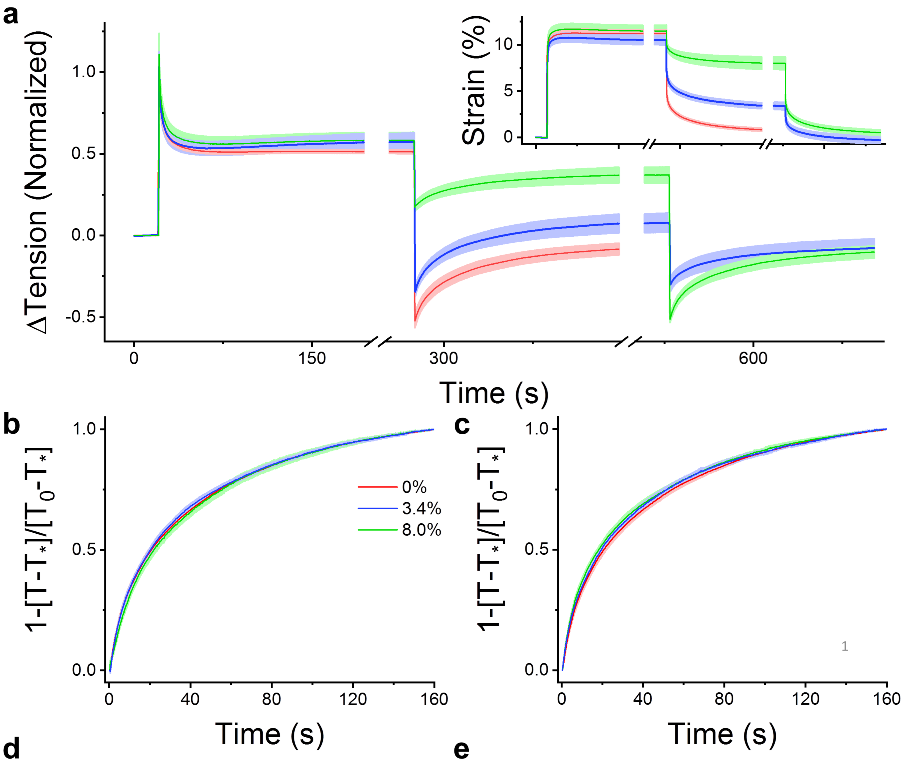


**d**

| | a (μN) | d (μN) | τ (s) | β |
|---|---|---|---|---|
| 0% | -5.6 ± 0.5 | -0.6 ± 0.3 | 42 ± 3 | 0.68 ± 0.02 |
| 3.4% | -5.3 ± 0.5 | 1.2 ± 0.6 | 40 ± 2 | 0.689 ± 0.008 |
| 8.0% | -2.6 ± 0.4 | 2.8 ± 0.5 | 39 ± 3 | 0.67 ± 0.03 |

**e**

| | a (μN) | d (μN) | τ (s) | β |
|---|---|---|---|---|
| 0% | -5.6 ± 0.5 | -0.6 ± 0.3 | 42 ± 3 | 0.68 ± 0.02 |
| 3.4% | -2.8 ± 0.3 | -0.7 ± 0.6 | 40 ± 3 | 0.67 ± 0.02 |
| 8.0% | -5.2 ± 0.5 | -0.8 ± 0.4 | 42 ± 4 | 0.64 ± 0.02 |

**SI 5: Microtissue stress recovery rate is strain independent.** Large step strains (insert) were applied and microtissues (N=6) were allowed to fully relax (a). In the next part of the experiment, microtissue recovery was assessed at a full return to initial length (red), and at large (blue) and small (green) intermediate steps. Recovery rates at 0%, 3.4% or 8.0% strain were identical (b). In the last part of the experiment, tissues at intermediary steps were returned to their initial lengths. These recoveries also shared the same rates (c). Panels (d) and (e) contain the average fitting constants to $-a * e\hat{}(-(t/\tau)\hat{}\beta) + d$ for the intermediate steps and the later recoveries to initial length, respectively. There were no significant differences in either the time or power law constants (repeated measures 1-way ANOVA; P>0.05) despite large changes to the step size and the strain at which the tissues recovered.

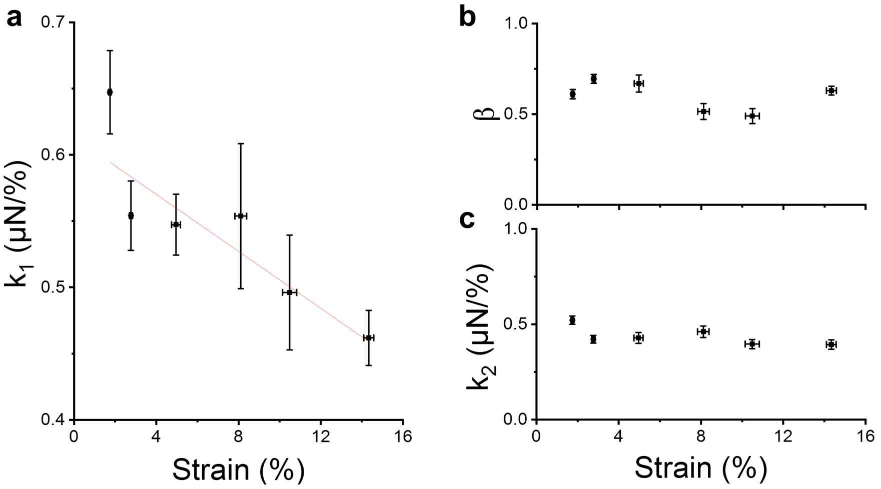


**SI 6: The fitting constants for microtissue stress relaxation at various step sizes.** To further explore nonlinearities, responses across 160 different microtissues were analyzed with step strains ranging from 1.75 ± 0.06% to 14.3 ± 0.2%. In agreement with repeated measurement findings presented in the body of the article, the microtissue $k_1$ spring constant, describing the amplitude of the relaxation response normalized to microtissue strain, linearly decreased with the step size (a) (linear regression; $R^2$=0.78 $P<0.001$). In contrast to repeated measurements, however, the β power law constant (b) and the $k_2$ spring constant (c) were invariant with step size ($P>0.05$). These conflicting results were not entirely surprising because the strain dependencies of these variables were small, and thus could be outweighed by inter-tissue variability. For example, at a given strain, the standard deviation of $k_2$ was on average 0.1 (or ~25% of the mean), which is considerable considering we only observed a 11 ± 5 % decrease between 3.3 ± 0.2 % and 10.9 ± 0.8% step strains. As for β, the standard deviation was equally as large in terms of its mean value (~24%) and the observed decrease with repeated measurements was also small (17 ± 4%) over the experimental range.

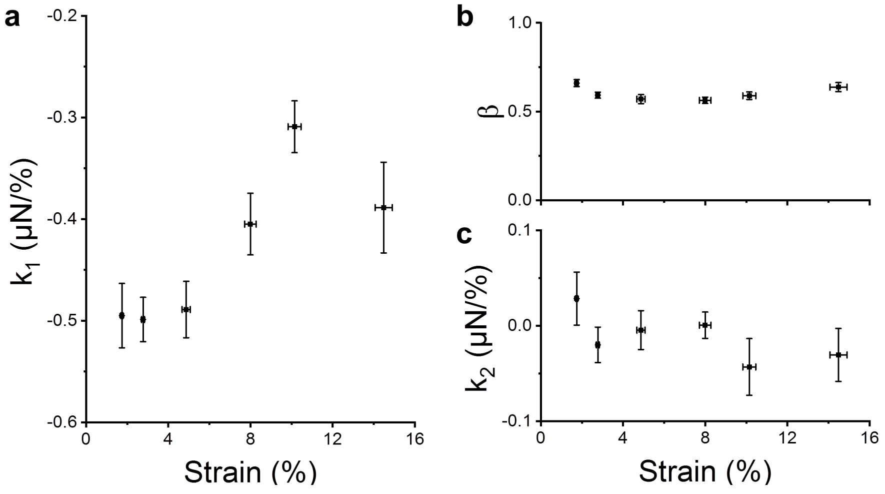


**SI 7: The fitting constants for microtissue stress recovery at various step sizes.** As in SI 5, the recovery fitting constants from 160 different microtissues are plotted against the step strain amplitude. In agreement with repeated measurement findings presented in the body of the article, the microtissue $k_1$ spring constant, describing the amplitude of the recovery response normalized to microtissue strain, decreased with the step size (linear regression; $R^2$=0.66 P<0.05). Further in agreement, the β power law constant and the $k_2$ spring constant were also invariant with step size. In addition, under all strain conditions, $k_2$ was approximately zero, indicating that there was no difference between residual stresses before and after the step strain experiment.

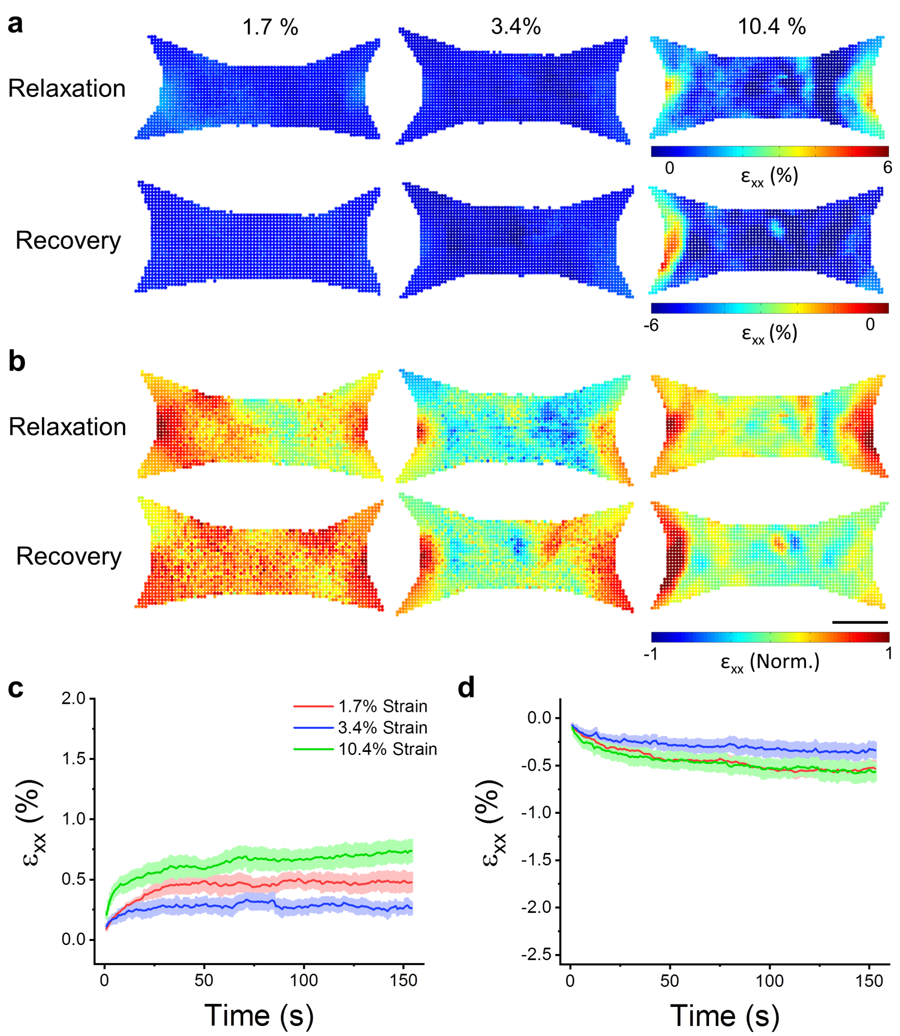


**SI 8: Remodeling in the longitudinal direction following changes to microtissue length.** Spatial distributions of relaxation and recovery in the longitudinal direction after various step sizes are shown in (a). The distributions are normalized to three standard deviations outside the absolute mean value in (b). The scale bar represents 100μm. The average (N=6) longitudinal strain during stress relaxation and recovery are shown in (c) and (d), respectively.

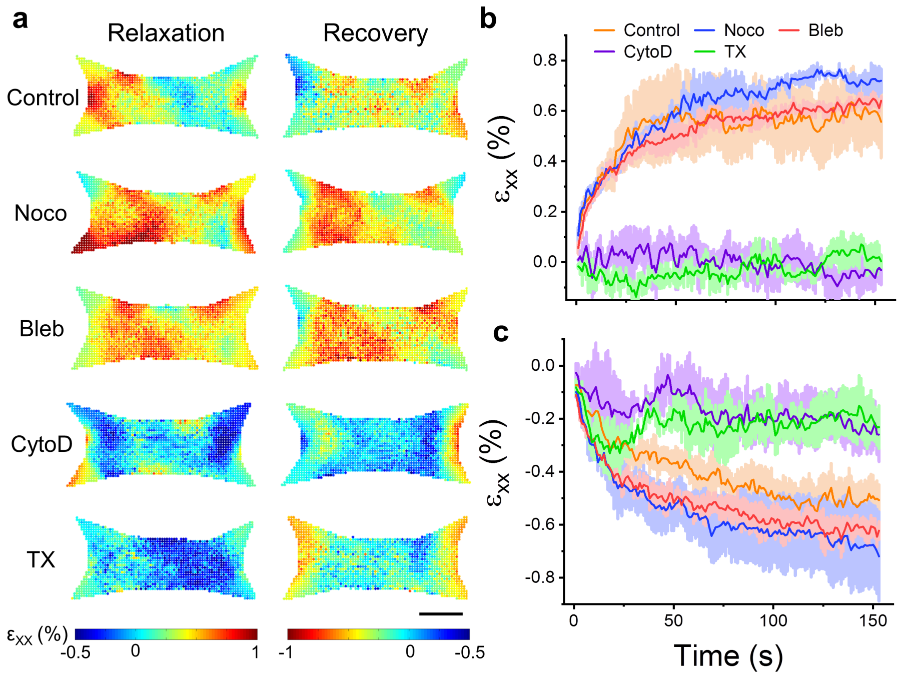


**SI 9: Remodeling in the longitudinal direction following changes to microtissues length varies with pharmacological treatments.** Spatial distributions of relaxation and recovery in the longitudinal direction after pharmacological treatments are shown in (a). The scale bar represents 100µm. The average (N=3) longitudinal strain during stress relaxation and recovery are shown in (b) and (c), respectively.
